# Taming the reference genome jungle: the refget sequence collection standard

**DOI:** 10.1101/2025.10.06.680641

**Authors:** Donald R Campbell, Timothee Cezard, Sveinung Gundersen, Andrew D. Yates, Robert M. Davies, John Marshall, Sang-Hoon Park, Alex H. Wagner, Michael I. Love, Reggan Thomas, Oliver Hofmann, Nathan C. Sheffield

## Abstract

Reference genomes are foundational to genomics but suffer from widespread ambiguity and incompatibility due to inconsistent naming, undocumented differences, and lack of formal mechanisms for comparison. To address this, we introduce the GA4GH refget Sequence Collections (seqcol) standard. Refget seqcol is a framework for unambiguous representation, retrieval, and comparison of sequence collections such as reference genomes and transcriptomes. The seqcol standard comprises four components: a structured data schema, a canonical encoding algorithm that produces content-based, globally unique identifiers, a retrieval API, and a comparison protocol. This standard enables precise identification of sequence collections, even across decentralized or private systems, and allows compatibility assessments beyond exact identity, such as order-relaxed matches or shared coordinate systems. We applied the refget seqcol standard to 60 human and 36 mouse reference genomes sourced from major providers. Using digest-based comparisons, we quantified levels of similarity across attributes including sequence names, lengths, coordinate systems, and actual sequence content. Our analysis revealed some consistent subsets of sequences or coordinate systems, as well as substantial incompatibility among references and duplicate references under different names. To support adoption of refget seqcol, we provide a Python package implementing the full standard, a web API, and a comparison interface allowing users to assess local references against a curated database. This work offers a scalable, reproducible solution to the reference genome compatibility crisis, enabling improved transparency, reuse, and integration in genomic analyses. Refget seqcol enhances interoperability across tools and datasets, making genomic research more robust and reproducible.

## Introduction

Reference genomes are used extensively in genomics. However, their broad reuse is complicated by the many different sources and versions of the same reference genome [1, 2]. Unfortunately, convenient human-readable names for reference genomes, such as “hg38” or “GRCh38,” do not unambiguously specify the source or version [3]. When reference genome data is not clearly specified or mismatches occur between analyses, the results may become incompatible or impossible to reproduce [4, 5]. This lack of clarity not only disrupts reproducibility but also impairs compatibility of downstream analysis results [6, 7].

However, not all discrepancies preclude interoperability. For example, results based on different references can be integrated when differences involve only non-essential chromosomes, and some analyses only require consistency of coordinate systems the set of sequence names and their corresponding lengths – without regard to sequences themselves [7]. Still, no formal tools or standards exist to help researchers assess whether two reference genomes or their derived data are compatible. This absence hinders the robustness and reproducibility of genomic research.

One solution is to unambiguously identify assemblies using unique accession keys, such as those from the NCBI Assembly database [8]. This addresses some concerns, but it relies on a central authority, limiting its use for private or custom genomes, and it does not accommodate multiple genome providers. Furthermore, centralized unique identifiers alone cannot *validate* identity, which also depends on sequence content. The previously published refget Sequences standard [9] addressed this by defining content-derived identifiers and a retrieval service for individual sequences, such as a single chromosome. The refget Sequences standard is now used widely, for example, as the CRAM reference registry, but it does not extend to collections of sequences like full reference genomes. To fill this gap, the refgenie reference genome manager [10] introduced unique, content-derived identifiers for sequence collections [11]. This approach catalyzed a broader community-driven effort to define use cases and develop a universal approach for identifying reference genomes.

Here, we present the refget Sequence Collections standard (“refget seqcol” for short). Developed through the Global Alliance for Genomics and Health (GA4GH) [12], the refget seqcol standard provides four components: a data schema; an algorithm for deterministic, computable identifiers; a retrieval API; and a comparison protocol. It is designed for reference sequences, including genomes and transcriptomes, but is applicable to any collection of sequences. We present several implementations of the standard, including a web server that allows users to compare a custom reference genome with a database of common reference genomes to assess compatibility for downstream analysis. To demonstrate how the standard improves interoperability, we quantified differences in coordinate systems and sequence content for 60 human reference genomes and 36 mouse reference genomes. Our analysis provides a comprehensive comparison of common references in use today, illuminating differences that might surprise some readers. It also demonstrates how refget seqcol can be used to assess similarities and compatibility among references, such as establishing shared coordinate systems, order-relaxed identity, name-relaxed identity, subset relationships, and more.

## Methods

### Components of the Sequence Collection standard

#### Overview of the standard

Refget Sequence Collections specifies several primary components (Fig. 1A): 1) a base schema that describes the structure of a set of input sequences; 2) an algorithm for generating deterministic identifiers from sequence collections themselves; 3) an API defining how to list, retrieve, and compare sequence collections.

**Figure 1:**
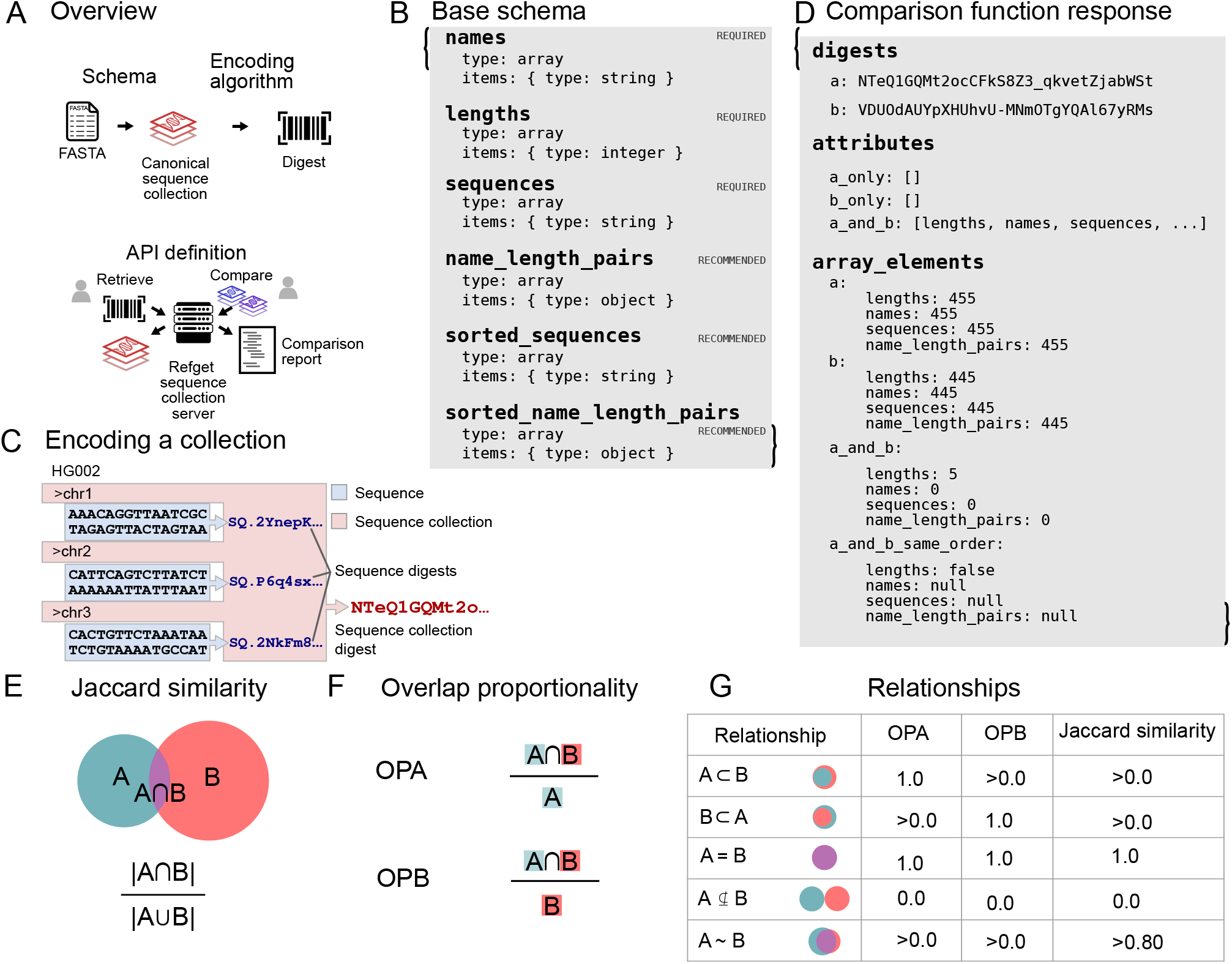
Overview of the refget Sequence Collections standard. A) Graphical overview showing how the four components of the standard fit together. B) Minimal “base” schema defining the data model for sequence collections. C) Graphical depiction of the encoding algorithm. D) Example result of a comparison between two sequence collections. E) Graphical depiction of Jaccard similarity, calculated as the intersection over the union for two sets. F) Graphical depiction of overlap proportionality, calculated as the intersection of two sets divided by the size of either of the sets. G) Table describing the inferred relationship of Set A to Set B depending on OPA/OPB and Jaccard similarity

##### 1. Base schema

Sequence collections are organized as an object with 3 required properties: names, lengths, and sequences (Fig. 1B). Each property is an array, with number of elements equal to the number of sequences in the collection, and with a shared index (these we call *collated* arrays). The data model may be customized with further attributes of two types: *inherent* attributes are those that contribute to the identity of the sequence collection. These attributes will be hashed as part of the encoding algorithm. In contrast, *non-inherent* attributes allow implementations to store and serve attributes that do not contribute to the digest. These non-inherent attributes enable identifiers to remain compatible even if servers differ in attributes offered. Each refget seqcol service specifies its data model by providing a schema that defines any attributes of the collection object and whether they are inherent. The standard recommends adding several non-inherent computable attributes, specifically `name_length_pairs, sorted_name_length_pairs` and `sorted_sequences`. These non-inherent attributes do not contribute to the identity of the sequence collection during hashing, but offer convenience for downstream use cases.

##### 2. Encoding algorithm

The encoding function specifies an algorithm that computes a unique digest for a set of named sequences (Fig. 1C). This digest acts as a universal, data-derived identifier. By basing the identifier on the data itself rather than relying on external naming conventions or authorities, this approach ensures any two systems using the same input sequences will compute the same digest, enabling consistent and unambiguous identification across different sources, including private or custom genomes [13]. Importantly, this method supports reproducibility in distributed or federated computing environments, even without shared naming standards. This function is generally provided by local software operating on a local set of sequences. The algorithm is described in detail in the formal specification (https://ga4gh.github.io/refget); briefly, the steps of the encoding process are:

- Step 1. Organize the sequence collection data into *canonical seqcol object representation*, which consists of vectors of *names, lengths*, and *sequences* (which are refget sequence digests).
- Step 2. Apply the RFC-8785 JSON Canonicalization Scheme (JCS) to canonicalize the value associated with each *inherent* attribute individually.
- Step 3. Digest each canonicalized attribute value using the GA4GH digest algorithm [14].
- Step 4. Apply the RFC-8785 JSON Canonicalization Scheme again to canonicalize the JSON of the new seqcol object representation.
- Step 5. Digest the final canonical representation again.

The result of encoding is the “top-level” digest, which provides a unique fingerprint for each collection of sequences. This approach provides several desirable properties. Because of the RFC-8785 JSON canonicalization, resulting digests do not depend at all on whitespace or JSON attribute order. The layered approach, where the individual properties are digested first, and then digested as a collection, provides a way of filtering, discovering, and standardizing on the values of individual properties. These intermediate digests are useful independent of the top-level digest.The encoding algorithm is generally very fast because it is restricted to the metadata about the sequences, so it scales with the number of sequences rather than their length. For typical collections with dozens to thousands of sequences, the computation completes in seconds.

##### 3. Retrieval and Comparison API

The standard specifies an API with these main endpoints, with addition sub-endpoints described in the specification:

1. `/service-info`, for describing information about the service.
2. `/collection`, for retrieving sequence collections.
3. `/comparison`, for comparing two sequence collections.
4. `/list`, for retrieving a list of objects.
5. `/attribute`, for retrieving the value of a specific attribute.

The primary utility of the retrieval API is to provide a lookup service that allows users to retrieve the exact sequence collection represented by a digest. These endpoints also allow users to discover collections on a server, filter them by attribute digests, or retrieve values of specific attributes. Importantly, the /comparison endpoint provides a way to check compatibility between two sequence collections that are not identical. Given two collections, this function describes the shared attributes, the number of shared and unique elements for each attribute, and whether the shared elements are in the same order (Fig 1D). This output provides enough information to establish various levels of compatibility, such as order-relaxed identity, matching sequences with different names, and subset relationships. This provides a basis for determining if results based on two different reference genomes can be integrated. One limitation of the comparison protocol is that it is restricted to attributes of the collection and does not extend to the level of individual sequence differences – any single nucleotide change results in a completely different sequence digest, and the comparison protocol only assesses whether the digests match.

### Analysis comparing common reference genomes

#### Data sources

To demonstrate the utility of the standard, we sought to explore the similarities and differences among commonly used reference assemblies. We first collected 68 human and 36 mouse reference genomes commonly used for sequencing analysis from a variety of different data distributors (Table 1, S1-S6). We included primary assemblies, patches, and variations containing decoy sequences.

**Table 1:** List of human and mouse reference providers and counts of included genomes.

| Provider | Human | Mouse |
| --- | --- | --- |
| 1000 genomes | 3 |  |
| Broad | 3 |  |
| DDBJ | 4 |  |
| ENA | 2 | 3 |
| Ensembl | 9 | 8 |
| GENCODE | 6 | 3 |
| iGenomes | 9 | 7 |
| NCBI | 17 | 6 |
| refgenie | 4 | 2 |
| UCSC | 11 | 7 |

#### Analysis

We downloaded FASTA files, processed them using the refget Sequence Collections encoding algorithm, and then applied the comparison function to compare each pair of genomes. We compared four attributes: names, lengths, sequences, and name-length-pairs. The *sequences* attribute consists of raw refget sequence digests. The name-length-pairs attribute concatenates name and length attributes and considers them as a unit, providing an assessment of coordinate systems without concrete sequences. To assess similarities and differences, for each attribute, we computed: 1) the Jaccard similarity (Figure 1E), providing an overall symmetrical assessment; and 2) OPA and OPB scores, which provide an asymmetrical overlap proportion with respect to either genome independently (Figure 1F). Together, these scores allowed us to define the relationships, such as subset/superset, between two reference genomes for each attribute (Figure 1G).

## Results

### Global Comparisons

#### Initial comparison of the 6 major providers

To understand high-level differences among major reference providers, we analyzed a limited set of 6 human hg38 references from six major providers: Ensembl, ENA, DDBJ, GENCODE, NCBI, and UCSC (Supplemental Table 1). For Jaccard similarity, 4 of 6 references have high pairwise similarities for sequences and lengths, indicating broad general agreement of the underlying sequences (Figure 2A). Compared to these four, the DDBJ reference had medium similarity for sequences and lengths and the ENA had low similarity. 5 out of 6 references had low similarity for names which, in turn, leads to low similarity scores for name-length pairs. The exception is the DDBJ reference which showed medium similarity for names and name-length pairs to the UCSC reference.

**Figure 2:**
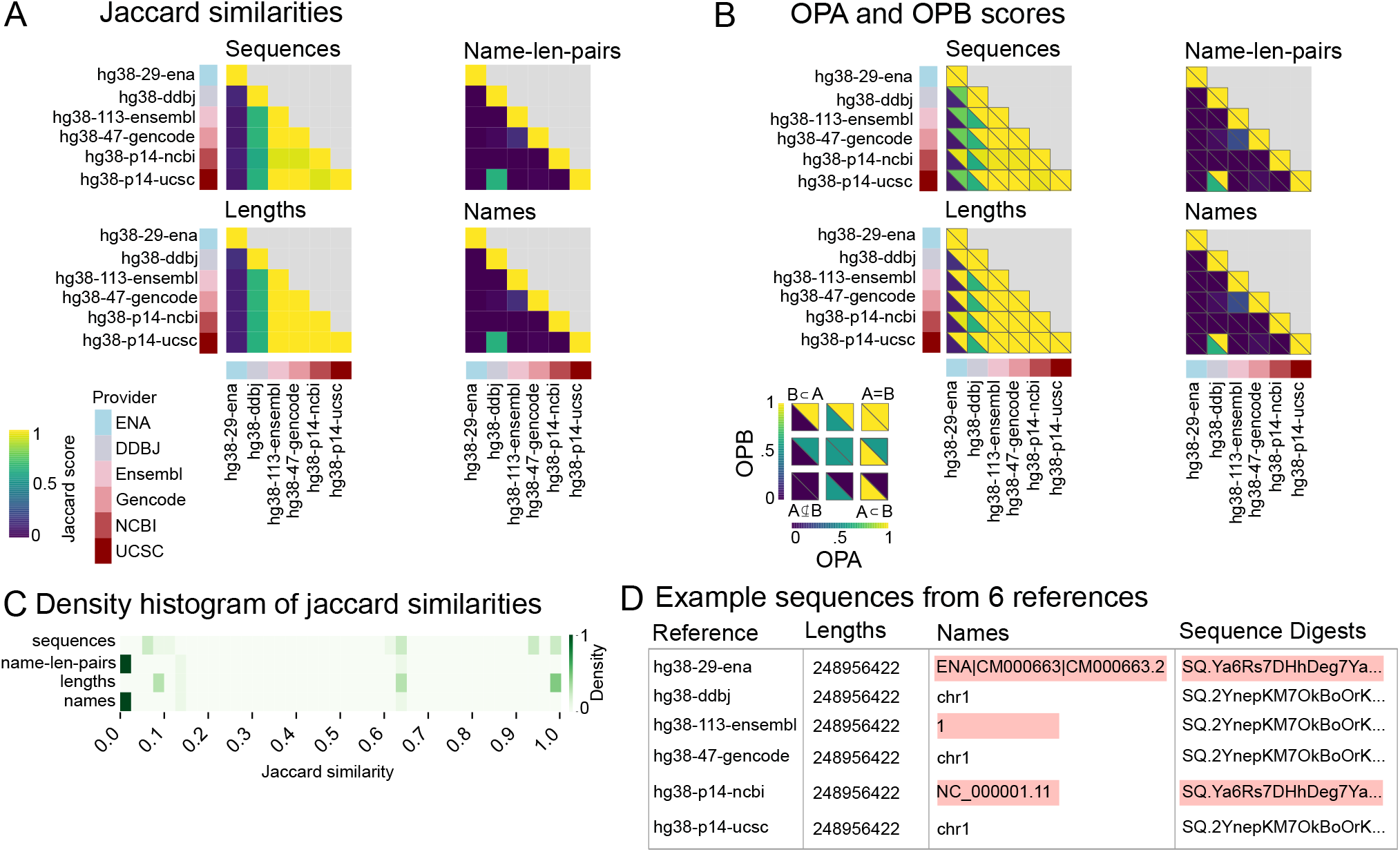
Results of comparison of primary human assemblies across 6 major providers. A) Jaccard similarity scores for sequences, name-length-pairs, lengths, and names. B) OPA and OPB scores for sequences, name-length-pairs, lengths, and names. C) Density map of Jaccard similarity scores for 15 pairwise comparisons of the 6 major providers. D) Diagram showing differences highlighted in red among 6 example reference genomes’ chromosome

To shed further light on the less-similar relationships, we compared the overlap proportionality scores (Figure 2B). High Jaccard scores imply high OPA and OPB scores, as illustrated in the group of 4 references with highly similar sequences and lengths. The ENA reference, which showed low similarity for sequences and lengths, has a high OPB score compared to the group of 4, indicating that this reference is a subset of the other references with regards to lengths, but its sequences are less similar. The DDBJ reference has a similar, but less pronounced relationship with the other 5 references. For names and name-length-pairs, there is little subsetting; the exception is the DDBJ and UCSC comparison flagged in the Jaccard score; the OP scores show that the DDBJ coordinate system is a subset of UCSC’s.

To summarize, we visualized a histogram map of Jaccard similarities for each attribute (Figure 2C). Most Jaccard similarity scores for names and name-length-pairs were close to 0, indicating an utter lack of agreement of sequence names among the providers. The situation is less bleak for sequences and lengths, for which we observed trimodal distributions, demonstrating some agreement among subsets of the providers for the underlying sequences. The raw data for chromosome 1 provides an anecdote of what gives rise to these results; each provider uses one of two underlying sequence variations and one of four name variations; the first chromosome in the ENA and NCBI databases has the same digest (raw sequences are identical), whereas the other four references use a different digest (Figure 2D). A manual check confirms that only two character changes are responsible for the sequence digest discrepancy.

Our analysis of mouse genomes for 5 major providers reveals high agreement for sequences and lengths for 4 out of the 5 providers (Supplemental Table 2, Supplemental Figure 1A). There is low similarity for sequence names and therefore for coordinate systems – the classic challenge of mismatched sequence names leading to incompatibility – with the exception of the one pairwise comparison between the GENCODE and Ensembl, which had an intermediate score. OPA and OPB scores show that none of these reference coordinate systems are subsets of each other (Supplemental Figure 1B). However, sequences and lengths are equal in most comparisons. Density map of Jaccard similarities shows most name and name-length-pair comparisons have low similarity, while the sequences and lengths show higher similarities than the human references (Supplemental Figure 1C).

#### Comparison of offerings from one major provider

Given the low level of similarity among the top reference genome providers, we were curious about concordance among versions of reference genomes from a single provider. Typically, providers issue derivatives of a genome with different alternative sequences or sequence processing, like masking. To compare these, we analyzed hg19 derivatives from UCSC (Supplemental Table 3, Figure 3).

**Figure 3:**
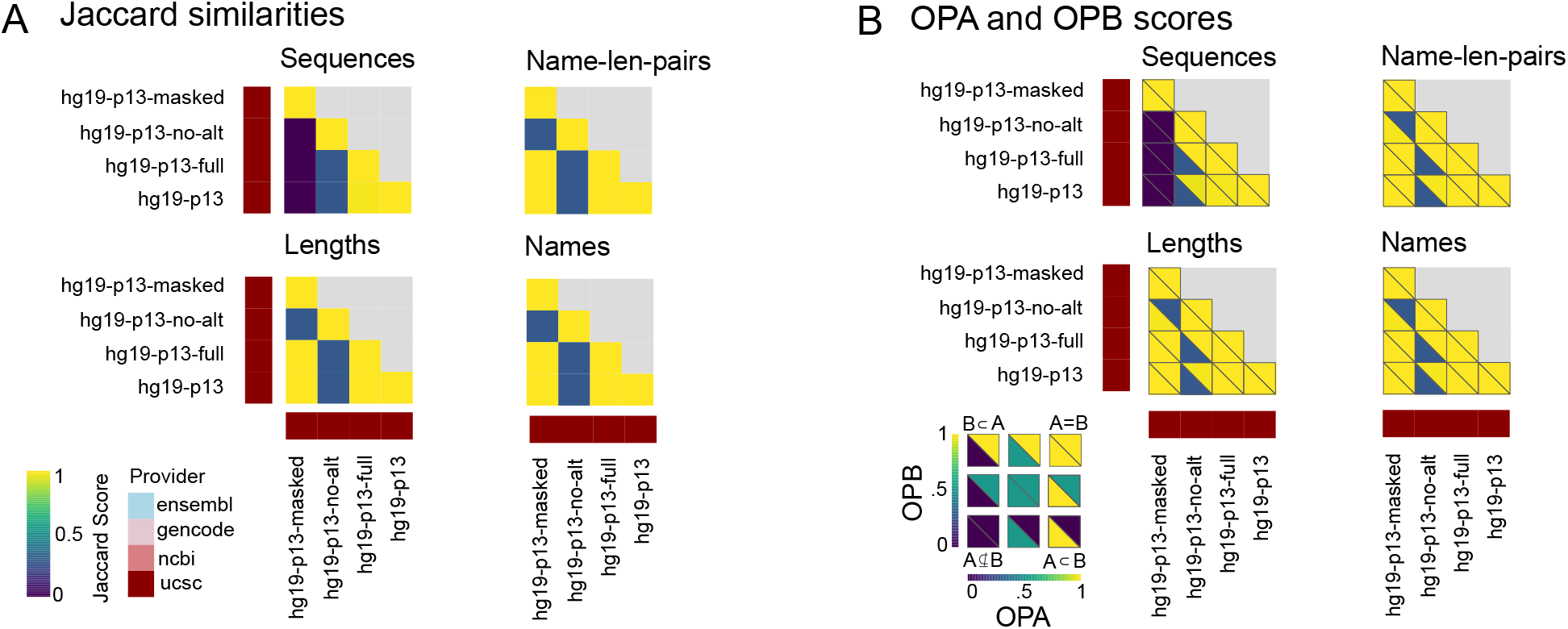
Comparing variations of UCSC’s hg19 patch 13. A) Jaccard similarity scores for sequences, name len, lengths, and names. B) OPA and OPB scores for sequences, name-length-pairs, lengths, and names.

Among these derivatives, the sequences were the least concordant, consistent with these derivatives frequently sharing names and lengths (Figure 3A). The noticeable outlier is the *hg19-p13no-alt* sequence, which has low Jaccard scores to all others for all attributes. As expected, the masked derivative differs from the others in sequence content. OPA and OPB calculations further reveal that the *no-alt* variant is a subset of all others, for all attributes except sequences, for which the masked derivative differs (Figure 3B). This indicates that, these derivatives all share a specific subset of a coordinate system, with matching names and lengths, which is the *no-alt* variant.

#### Comparison of a provider’s patches over time

Updates to the human reference commonly occur through patches, which modify the reference genome without disrupting the existing coordinate system [15]. To explore concordance among patches, we analyzed 9 patches of one reference genome from one provider (Supplemental Table 4), from the initial release of GRCh38 by NCBI (indicated by p0, from December 17, 2013) to the most recent patch, p14 (released on February 3, 2022). Our comparison illustrates that similarity decreases over time as the reference genome is patched and new versions are released (Figure 4A). OPB scores indicate that older patches are always subsets of newer patches for all four attributes, consistent with the constraint of patches to maintain backwards compatibility (Figure 4B).

**Figure 4:**
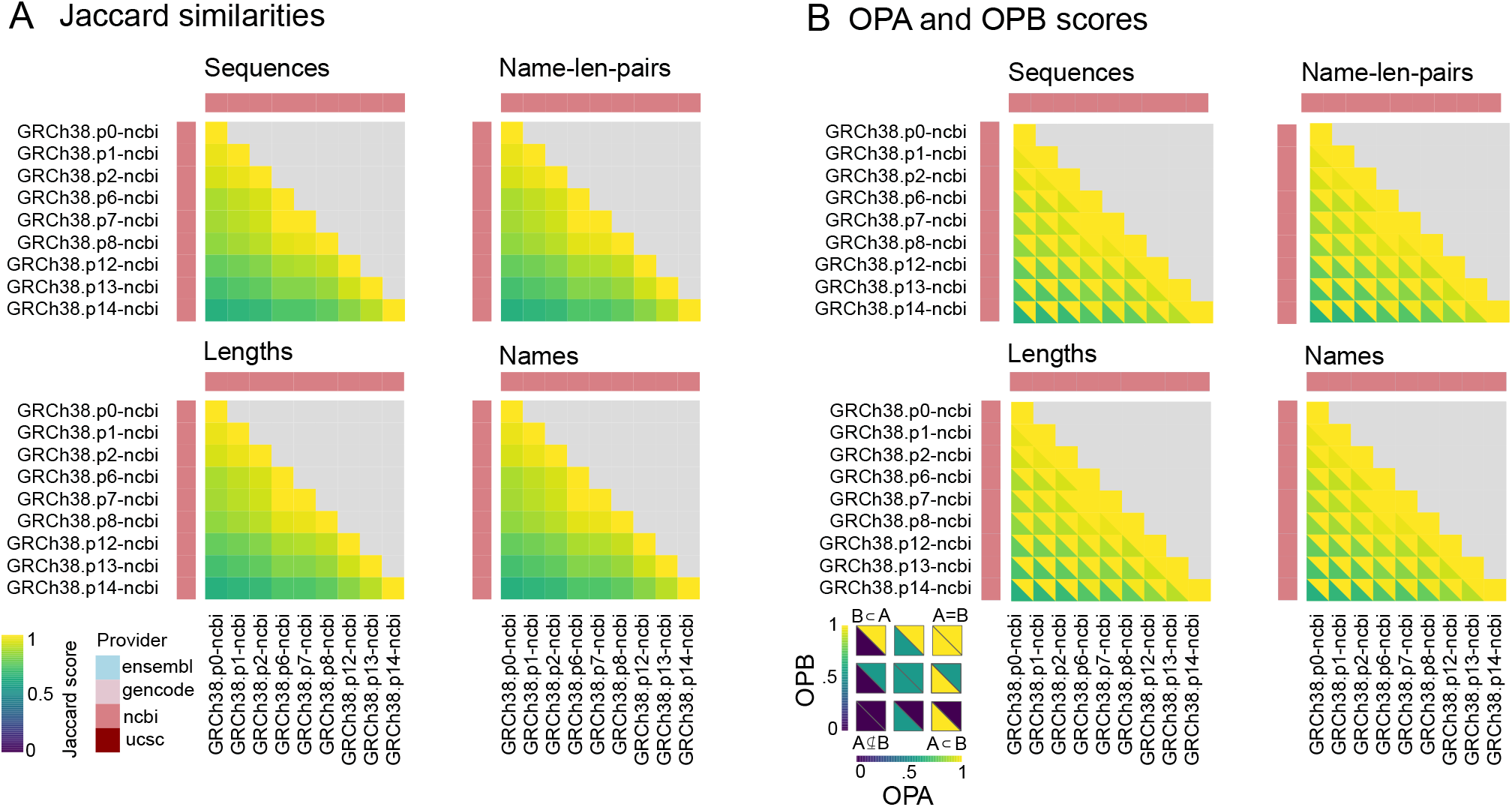
Comparing patches over time for NCBI GRCh38. A) Jaccard similarity scores for sequences, name-length-pairs, lengths, and names. B) OPA and OPB scores for sequences, name-length-pairs, lengths, and names.

#### Comparison of 60 human reference genomes and 36 mouse reference genomes

Having observed significant divergence between providers and even between *variations* from the same provider, we turned to analysis of all 60 human reference genomes (Supplemental Table 5). Scores were generally low, with sequences among the higher scores (Figure 5A), name-length pairs the lowest (Figure 5B), lengths the most similar (Figure 5C), and names the most disjoint of the 3 primary attributes (Figure 5D). Overall, there was little agreement between name-length-pairs and names and only limited agreement for lengths and sequences (Figure 5E). Comparing the providers, the ENA references showed the greatest difference across all attributes (Figure 5F). Other providers, such as GENCODE, showed a large difference in lengths and sequences but not names. Plotting the average jaccard score for each provider showed that lengths had the highest jaccard scores across providers, followed by sequences (Figure 5G). Out of the 60 human reference genomes, 28% were duplicates. For attributes, lengths had the greatest number of identical matches and sequences the least (Figure 5H). We also found lengths had the greatest and sequences had the least subsetting (Figure 5I).

**Figure 5:**
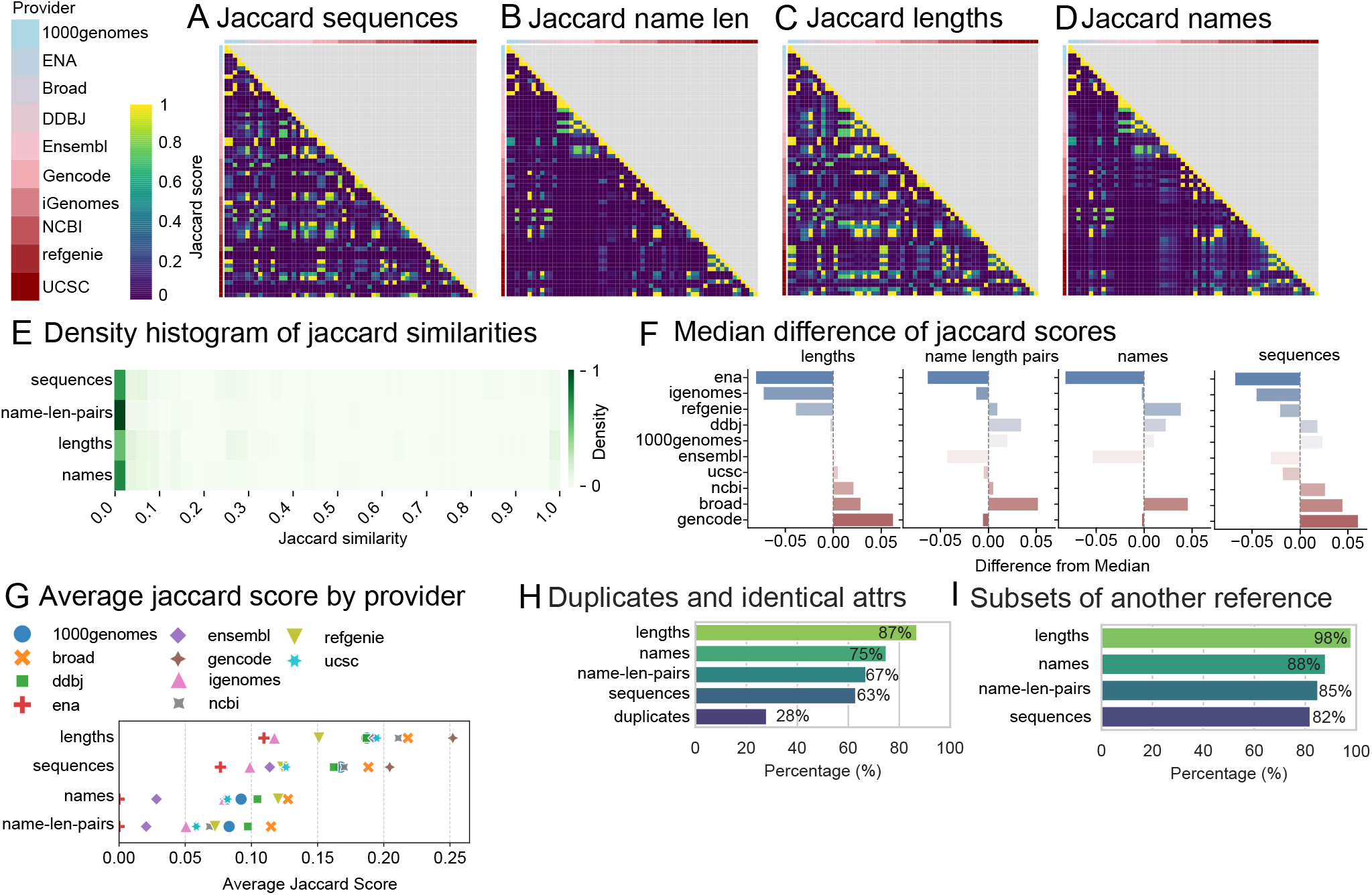
Comparing Jaccard similarities for 60 human reference genomes, grouped by provider. A) Jaccard similarity scores for sequences B) Jaccard similarity scores for name length pairs C) Jaccard similarity scores for lengths D) Jaccard similarity scores for names E) Density map of Jaccard similarity scores for 1770 pairwise comparisons. F) Median difference plots for each attribute grouped by provider G) Average jaccard scores for each provider H) Bar chart showing percentages of duplicates and identical references based on attributes. I) Bar chart showing percentage of references that are subsets of each other based on attributes.

Results for 36 mouse reference genomes (Supplemental Table 6), were similar, with sequences being second most similar (Supplemental Figure 2A) and name-length-pairs having the lowest scores (Supplemental Figure 2B). The lengths attribute had the highest scores (Supplemental Figure 2C), and the names attribute was third highest (Supplemental Figure 2D). Overall, there was little agreement between name-length-pairs; however, there is more agreement among sequences and lengths for mouse genomes when compared to human genomes (Supplemental Figure 2E). Among providers, iGenomes had the largest difference for the lengths and sequences attributes while ENA had the greatest difference for the names attributes and tied GENCODE for the name-length pairs attribute (Supplemental Figure 2F). Plotting the average Jaccard score for each provider showed a similar pattern to the human data (Supplemental Figure 2G). A notable exception is iGenomes where the names attribute scored highest.

Mouse genomes were duplicated less frequently (22%) than humans, but contained more identical attribute sets. The lengths attribute had the most identical sets while sequences had the fewest (Supplemental Figure 2H). Like in humans, the lengths attribute had the most subsets and sequences had the least. Mouse reference genomes had less subsetting when compared to human (Supplemental Figure 2I).

#### Comparison of sequence presence across all human references

We next investigated whether specific sequences were used frequently among many references. We divided the reference into major versions (hg38, hg19, *etc*.) and assessed the frequency of each sequence in the class. For hg38, 4775 independent sequences appeared in at least one reference. Most of these sequences were decoy sequences concentrated in few references (Figure 6A). Beyond these few references containing many sequences, there is a sharp drop off in the number of sequences across the remaining genomes. Similarly for hg19, 600 sequences appeared, with most reference genomes having few sequences (Figure 6B). However, the few references that contain many sequences share mitochondrial sequences. Finally, our dataset only included 3 hg18 references, with 2 of the 3 holding the mostly the same sequences (Figure 6C).

**Figure 6:**
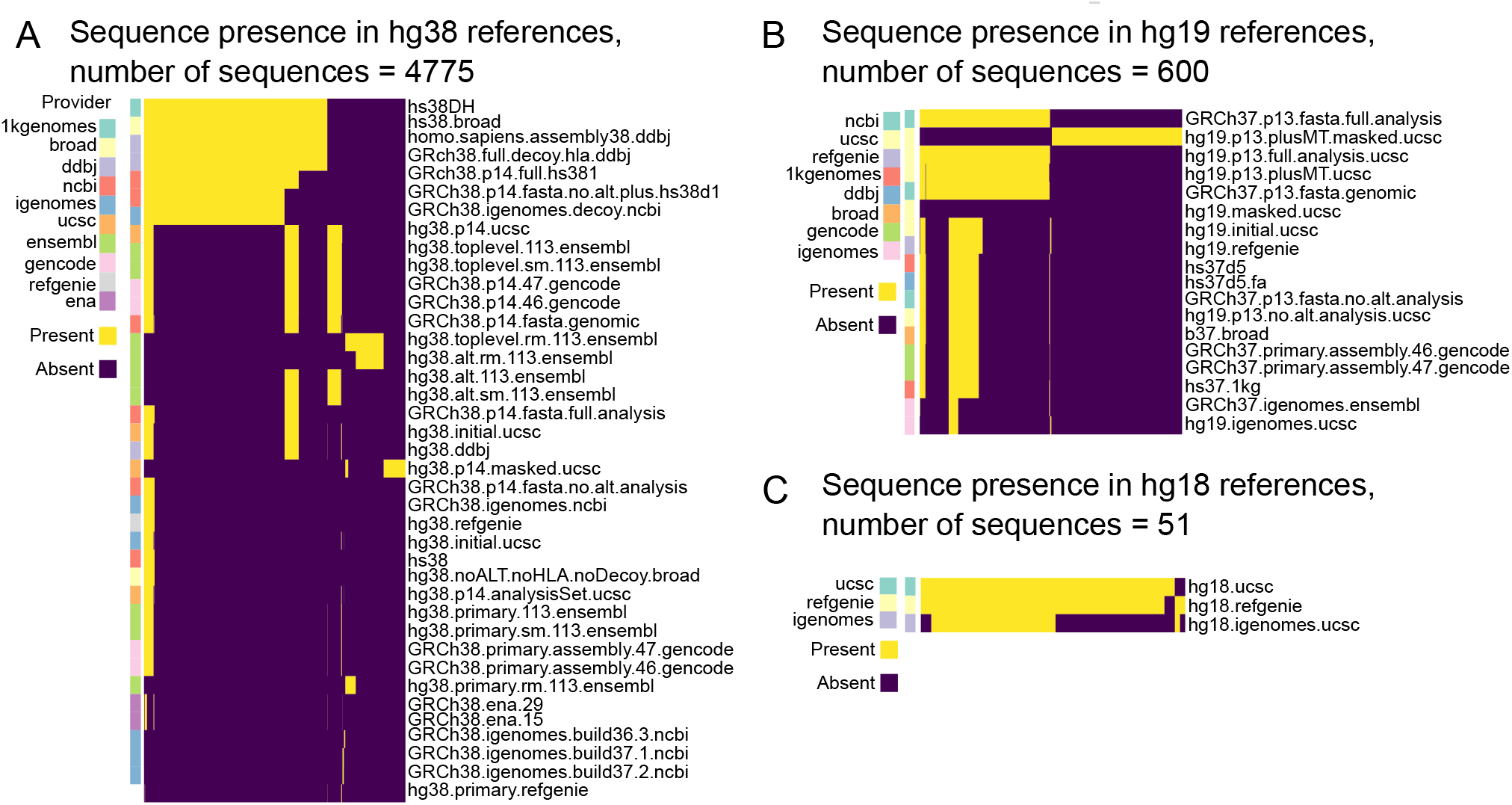
Comparing the presence of sequences within human references. A) Comparing 4775 sequences within hg38 references B) Comparing 600 sequences within hg19 references C) Comparing 51 sequences within hg18 references.

In mouse, the mm39 genomes had 607 sequences with top-level genomes from Ensembl and GENCODE having the largest presence (Supplemental Figure 3A). The mm10 genomes had 478 sequences with ucsc genomes dominating the counts with either soft or hard masked genomes (Supplemental Figure 3B). For m38, there are 239 sequences with the references procured directly from NCBI having the majority presence. Interestingly, the NCBI reference procured from iGenomes had a much smaller sequence presence (Supplemental Figure 3C). Our mm9 references had only 35 sequences, with over 50% being shared (Supplemental Figure 3D). Finally, m37 had the fewest sequences, all of which were shared among the analyzed references (Supplemental Figure 3E).

### Implementations and use cases

#### Existing and planned implementations

Several implementations of the refget seqcol standard are being developed. The reference implementation in the `refget` Python package includes three main components: (1) local digest computation tools, which a fast Rust implementation to compute seqcol identifiers from FASTA or similar inputs; a `SequenceCollectionClient` class, for querying remote seqcol-compliant APIs; and (3) a FastAPI-based server framework, which implements the full API specification and includes a `RefgetDBAgent`, which connects to a SQL backend for storing sequence collections. The package also includes a compliance test suite and data to validate against the API and local digest computation specifications. A second implementation is being developed by the European Variation Archive (EVA), featuring a Java-based server backed by a PostgreSQL database (https://github.com/EBIvariation/eva-seqcol). This implementation is built around the ability to dynamically retrieve and register reference genomes directly from INSDC, enabling broad and automatic support for all assemblies commonly used in genomic analysis. It also supports multiple existing naming conventions, including those used by ENA, GenBank, RefSeq, and UCSC, improving interoperability across diverse tools and datasets. By integrating with established reference sources and supporting common aliases, the EVA implementation reduces the friction of adopting sequence collections in existing workflows. An alternative client implementation is under development as part of the Omnipy Python package (https://omnipy.readthedocs.io/), with a focus on simplifying batch queries and other operations on sequence collections. Finally, the Ensembl project intends to deploy a public instance, based on our new beta infrastructure and metadata APIs, providing access to sequence collections across the tree of life including high profile projects such as Tree of Life and the Human Pangenome Reference Consortium. The first release of this service is due in early 2026 and will provide data as loaded into our beta infrastructure (https://beta.ensembl.org).

#### Web UI for comparing against a database of reference genomes

We have deployed a public reference implementation of the API at https://refget.databio.org, which supports several common use cases. For example, users can query the server to list all available sequence collection digests, or to retrieve a specific collection using its top-level digest. To check whether a local reference exists on the server, users can compute the digest of a local FASTA and search for a match. To demonstrate tools that can be built on top of the seqcol standard, we extended the server with several web applications to address common use cases. For example, one common use pattern we anticipate is to compare a local reference genome against genomes hosted on the server (Figure 7A). To make it simpler to interpret results of the comparison function, we developed the Seqcol Comparison Interpretation Module (SCIM), which allows a user to paste the output of a comparison endpoint and get a human-friendly interpretation of the result (Figure 7B; https://refget.databio.org/scim). This module interprets the numbers of the comparison into a simpler, understandable observation about how related two sequence collections are. To extend this reference discovery, or identifying existing collections that are similar in some way to a user-provided query, we developed the Seqcol Comparison Overview Module (SCOM), which goes beyond the 1-to-1 query provided by the comparison API to a 1-vs-many comparison. Users with a query collection can send this to the server, which iteratively compares it to each of the human or mouse reference genomes assembled for the analysis in this paper, and aggregates the results (Figure 7C; https://refget.databio.org/scom). The resulting summary figure describes similarity scores for the 4 key attributes, highlighting existing references that may be usable for the analysis, encouraging reuse (Figure 7D).

**Figure 7:**
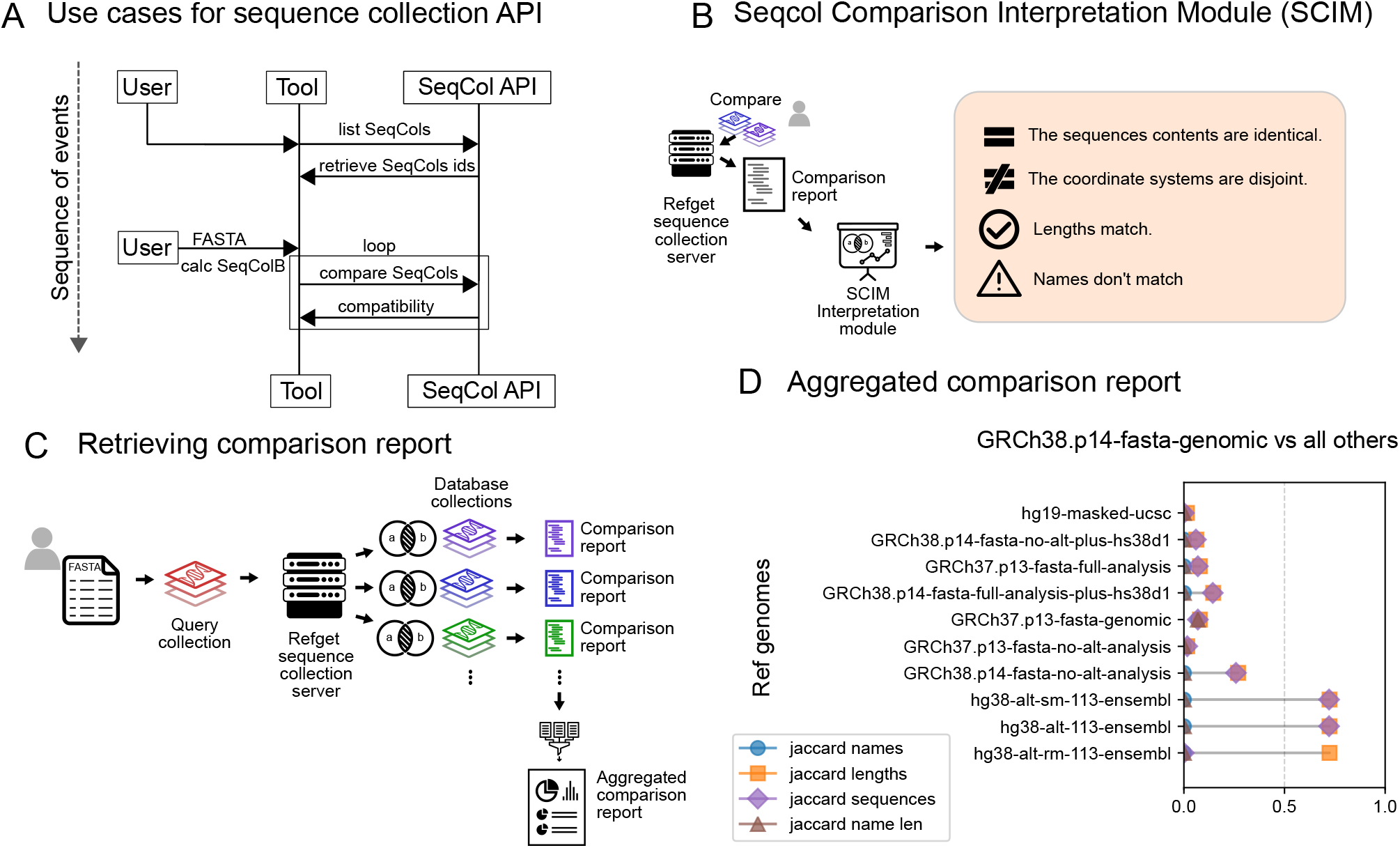
Implementations of refget sequence collections. A) Process diagram for usage of sequence collection API B) Diagram for comparison report retrieval C) Two example comparisons of one reference versus many. D) Example of an aggregated comparison report.

## Discussion

Sequence Collections describes a microservice for reference genome data [16] that solves many longstanding challenges with reference genomes. It provides: 1) globally unique, deterministic, computable identifiers; 2) a service for reference retrieval; 3) ability to uniquely identify coordinate systems or sequence collections without regard to sequence names; 4) a powerful comparison API that can assess shared coordinate systems, order-relaxed identity, name-relaxed identity, subset relationships, and other compatibility tests.

In terms of identification, the refget seqcol standard uses content-derived identifiers [9, 17], which provides several advantages over existing approaches. As a third-party standard independent of any specific reference provider, it does not rely on or create a central authority. Seqcol identifiers can be minted and verified by anyone, leading to automatic federation among all who use the standard, including data providers and private users. It also makes sequence content easily verifiable, as any adjustment to reference data content is reflected in the identifier, regardless of how it is referenced [7]. Because sequence collections identifiers are deterministic and content-derived, they can ensure that references from different locations are identical not only in name, but in content. Furthermore, publications that include the refget Sequence Collection digest of the reference genome used for the analysis will make it possible for downstream users to not just identify, but programmatically retrieve the exact sequence collection used in the analysis.

Beyond identification of entire genomes, seqcol also provides features for components of collections. For example, a coordinate system consists of a set of sequence names and their corresponding lengths, but without specific concrete sequences; it may be thought of as a sequence collection in which the sequences themselves are hidden or ambiguous. Coordinate systems provide an abstraction layer in which multiple sequence collections may subscribe to the same coordinate system despite differences in sequence content. Through the `name_length_pairs` attribute, seqcol provides a unique identifier for coordinate systems, adding functionality to coordinate-based annotations, which may generally be transferred from one sequence to another only if their coordinate systems match. The comparison endpoint automatically uses this identifier to compare whether annotations made on two different reference genomes may be compatible, despite differences in underlying sequences. This coordinate system identification also fills an unmet need in biodiversity sequencing projects [18], where annotation and visualization tools [19–21] can make use of the `sorted_name_length_pairs` identifiers for coordinate systems without respect to sequence order. This advantage matches the order-invariant coordinate system underlying an instance of a genome browser and other tools such as colocalization analyses [22, 23].

To our knowledge, our analysis of reference genomes is the most comprehensive comparison to date among available human and mouse references, and sheds light on the current state of reference compatibility. The results demonstrate an overwhelming lack of identity among sequences, lengths, names, and name-length-pairs across most human and mouse references. This result highlights and quantifies the widely known prevalence of reference genome variation. Though commonly ignored, this is a major problem in genomics, and it is particularly problematic for integrating data in secondary analysis. On the positive side, we also observed meaningful pockets of identity, including many subset relationships or other partial compatibility, such as consistent subsets of sequences, even if names differ. Furthermore, nearly a third of references were duplicates masquerading under different names, and many more were subset of another reference. Through the /comparison endpoint, refget sequence collections allowed us to identify these subsets, going beyond the simple comparison of identity that can be done by comparing the top level digest, and providing a more quantitative assessment of compatibility than simply ending with “not the same,” as is so common in analysis. However, this comparison approach is limited in its reliance on Jaccard similarity and overlap proportions, which reduce the complexity of sequence similarity to simple binary measures. These measures prioritize identity over closeness, as the standard does not support comparisons at the level of individual sequence differences – any single nucleotide change results in a completely different sequence digest. While this strict identity criterion ensures unambiguous equivalence and is therefore beneficial for reproducibility and reusability, it does not enable recognition of near-identical sequences. As a result, our findings of widespread divergence among sequences obscure substantial underlying similarity and large regions of congruence across the reference assemblies.

With the standard now approved by GA4GH, seqcol is poised to make analyses more reproducible and simplify integration of annotations from different sources. We are working to integrate seqcol across the genomics ecosystem; for example, the GA4GH Variant Representation Specification (VRS) can use refget seqcol to define the reference on which a variant is based [17]. Similarly, the GA4GH Data Repository Service (DRS) could extend a seqcol server to provide reference FASTA file downloads using seqcol identifiers. Seqcol identifiers can strengthen reference provenance by being embedded into file formats like SAM/BAM or VCF [24], in metadata headers like FAIR BioHeaders for Reference Genomes [5], or in genomic annotation standards developed in the Research Data Alliance (https://www.rd-alliance.org/groups/fairification-genomic-annotations-wg). For circular genomes, a SEGUID v2 digest can be combined with refget seqcol to derive identifiers for collections of circular genomes [25]. Toolchains such as salmon/tximeta [26] that use computed identifiers for automated transcriptome annotation intend to update to the seqcol identifiers in order to support additional use cases where only names and lengths, but not full reference sequences are available. The refget seqcol framework may also be extended to support pangenomes and graph-based references [27–29]. Considering a pangenome as collections of linear references, an extended refget seqcol that supports collections of sequence collections will enable similar deterministic, content-derived identifiers of pangenomes. This would provide benefits of precision, traceability, and interoperability for pangenome references, enabling consistent identification and comparison of increasingly complex reference models across tools and datasets.

## Acknowledgements

We are thankful for helpful discussions in the GA4GH.

## Data availability

The formal specification is hosted at: https://ga4gh.github.io/refget/

The reference implementation can be found at: https://github.com/refgenie/refget

All calculated results for human references can be found in Supplemental File 1 and all results for mouse references can be found in Supplemental File 2. Human reference sequence presence results can be found in Supplemental File 3. Mouse reference sequence presence results can be found in Supplemental File 4.

## Supplemental Figures

**Supplemental Figure 1:**
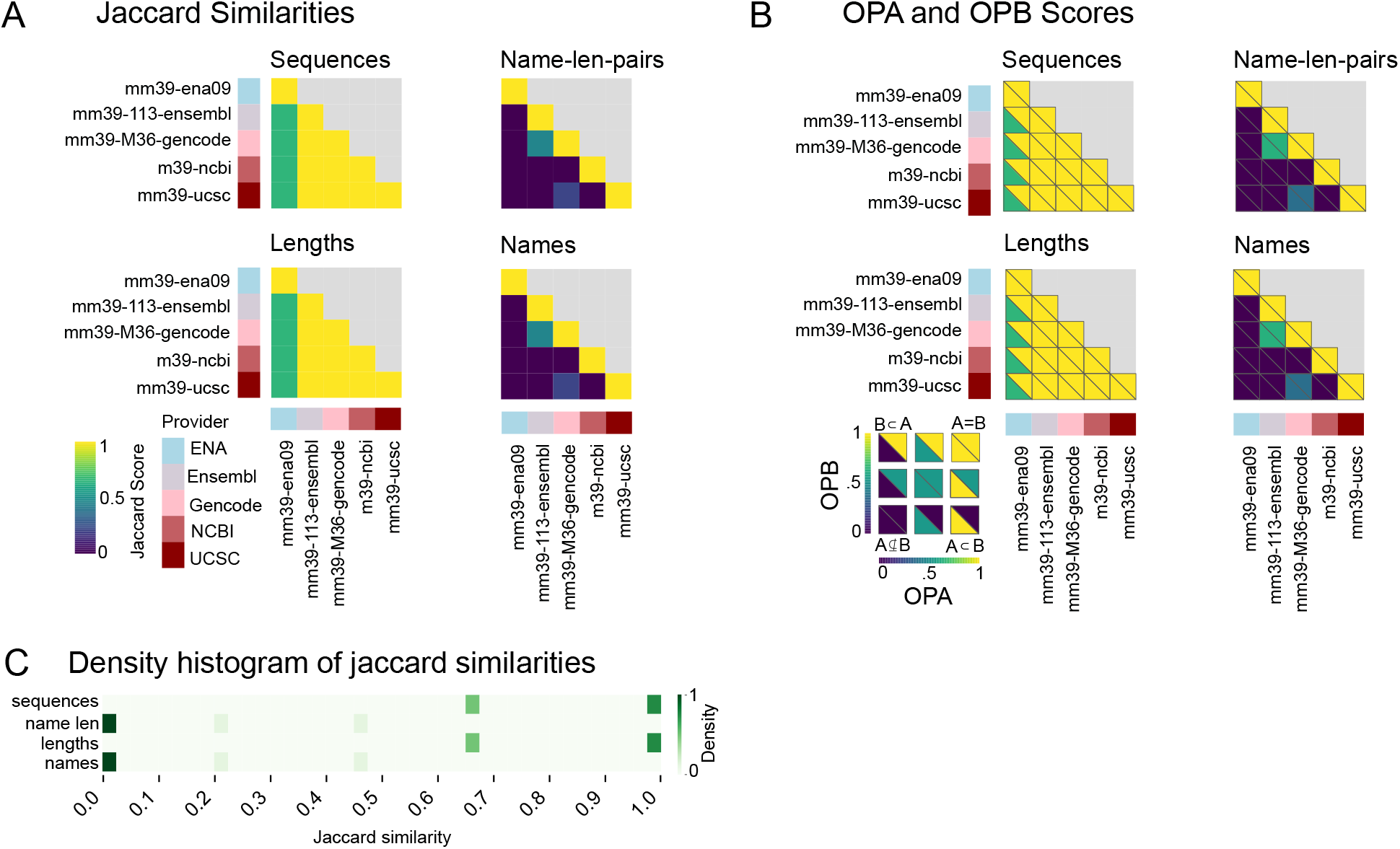
Results of comparison of primary mouse assemblies across 5 major providers. A) Jaccard similarity scores for sequences, name-length-pairs, lengths, and names. B) OPA and OPB scores for sequences, name-length-pairs, lengths, and names. C) Density map of Jaccard similarity scores for 10 pairwise comparisons of the 5 major providers.

**Supplemental Figure 2:**
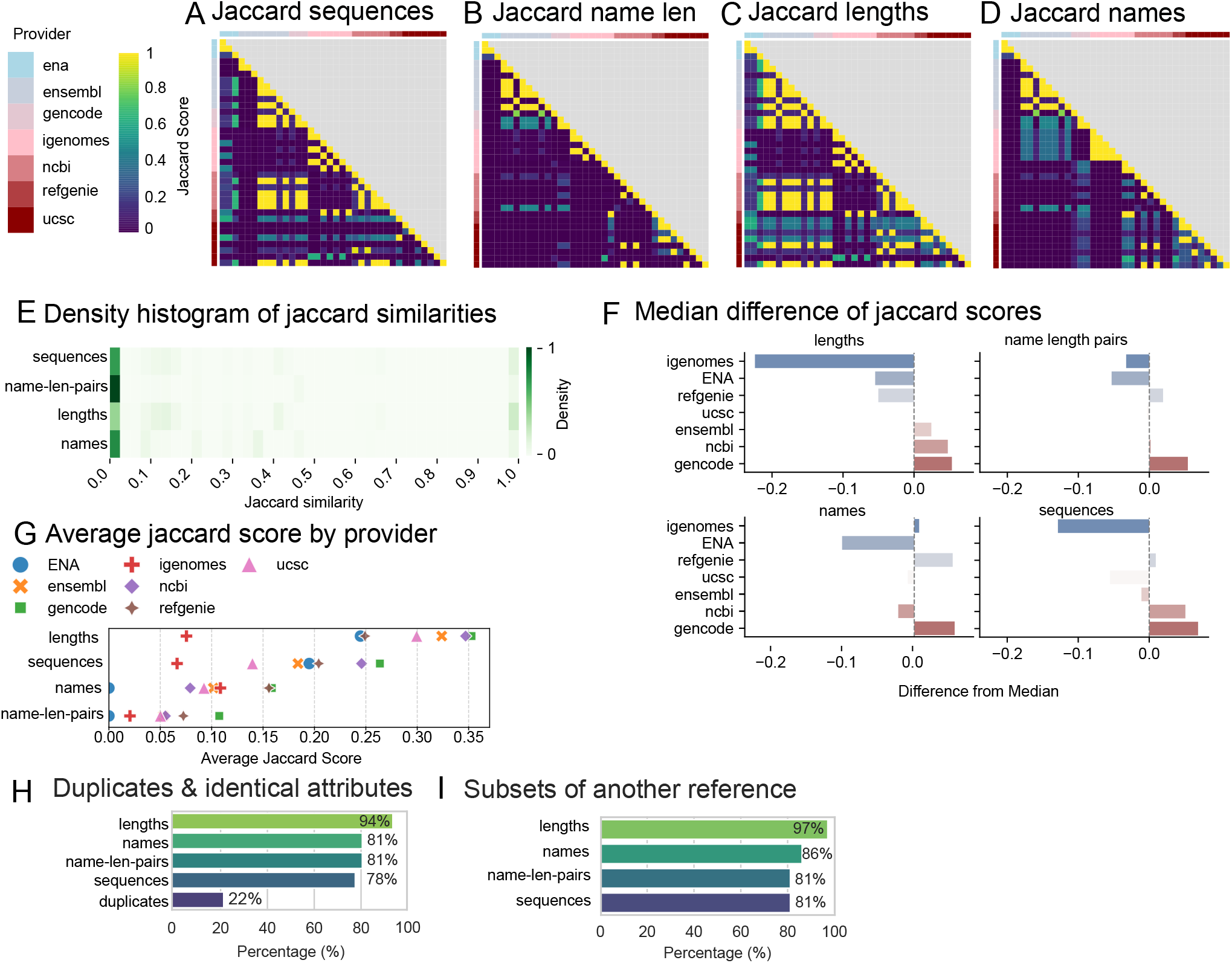
Comparing Jaccard similarities for 36 mouse reference genomes, grouped by provider. A) Jaccard similarity scores for sequences. B) Jaccard similarity scores for name length pairs C) Jaccard similarity scores for lengths. D) Jaccard similarity scores for names. E) Density map of Jaccard similarity scores for 630 pairwise comparisons. F) Median difference plots for each attribute grouped by provider G) Average jaccard scores for each provider H) Bar chart showing percentages of duplicates and identical references based on attributes. I) Bar chart showing percentage of references that are subsets of each other based on attributes.

**Supplemental Figure 3:**
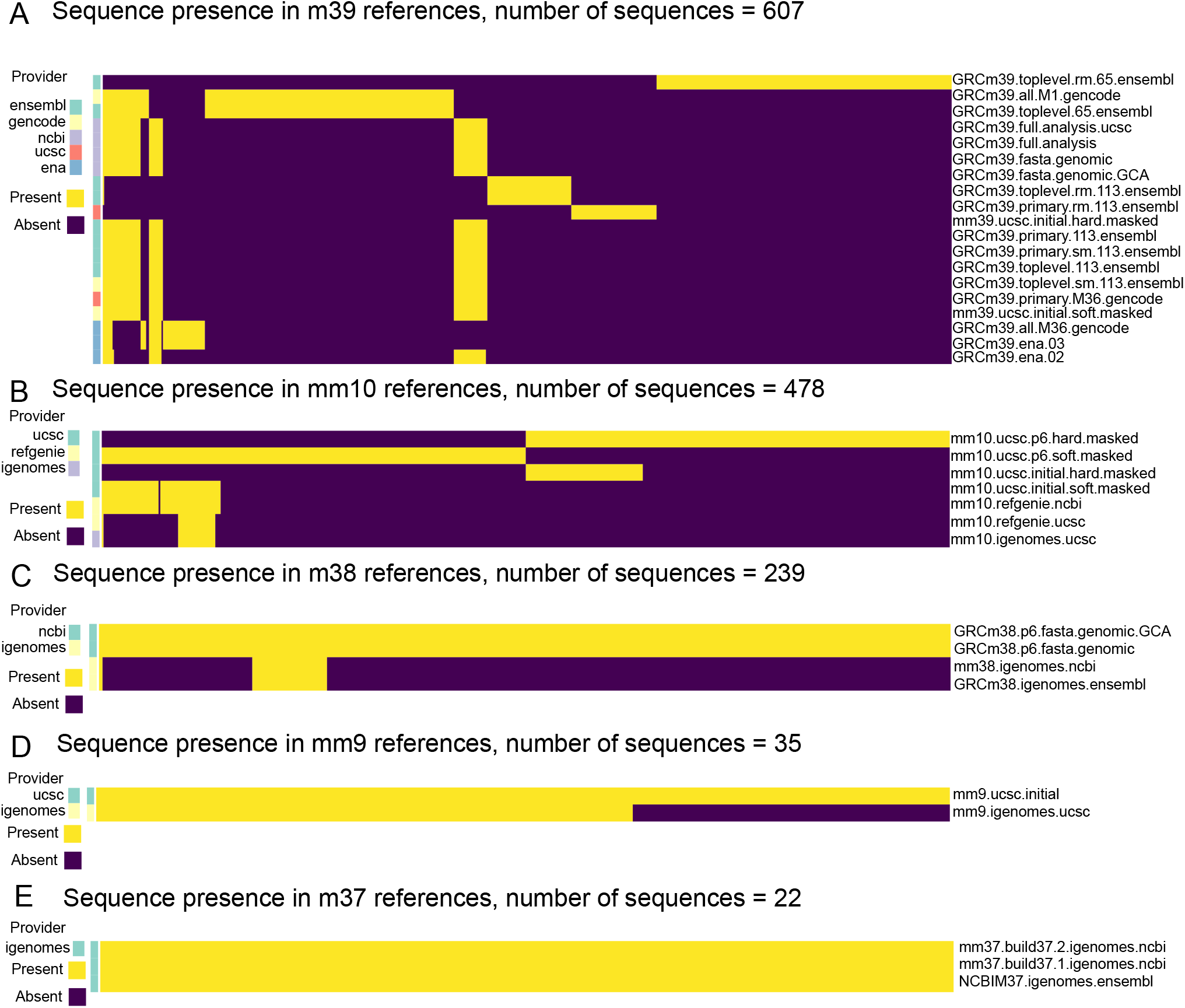
Comparing the presence of sequences within mouse references. A) Comparing 607 sequences within m39 references B) Comparing 478 sequences within mm10 references C) Comparing 239 sequences within m38 references D) Comparing 35 sequences within mm9 references E) Comparing 22 sequences within m37 references.

## Supplemental Tables

**Supplemental Table S1:**
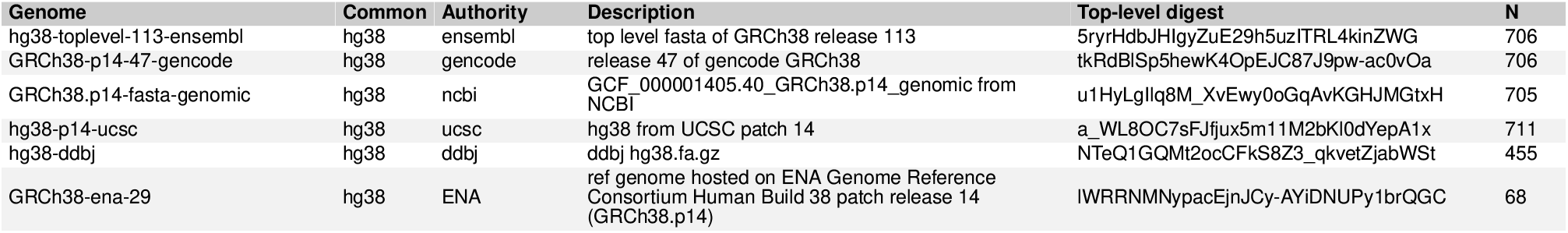
6 major human reference providers. This table shows major providers used for the core analysis. N=number of sequences present in the reference genome.

**Supplemental Table S2:**
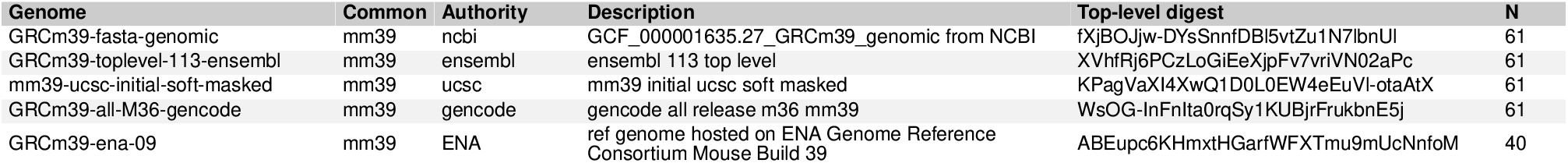
5 major mouse reference providers. This table shows major providers used for the core analysis. N=number of sequences present in the reference genome.

**Supplemental Table S3:**
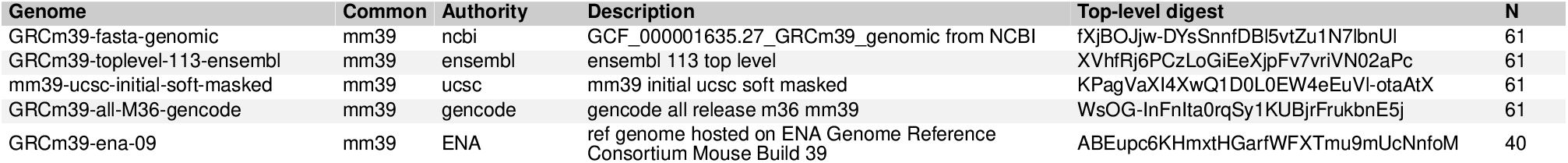
Selection of hg19 patch 13 references. N=number of sequences present in the reference genome.

**Supplemental Table S4:**
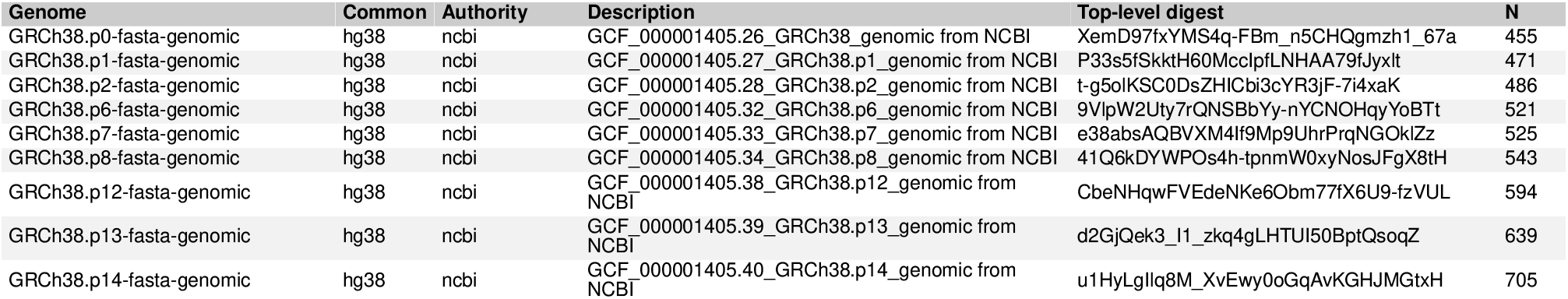
Selection of human NCBI GRCh38 patches. N=number of sequences present in the reference genome.

**Supplemental Table S5:**
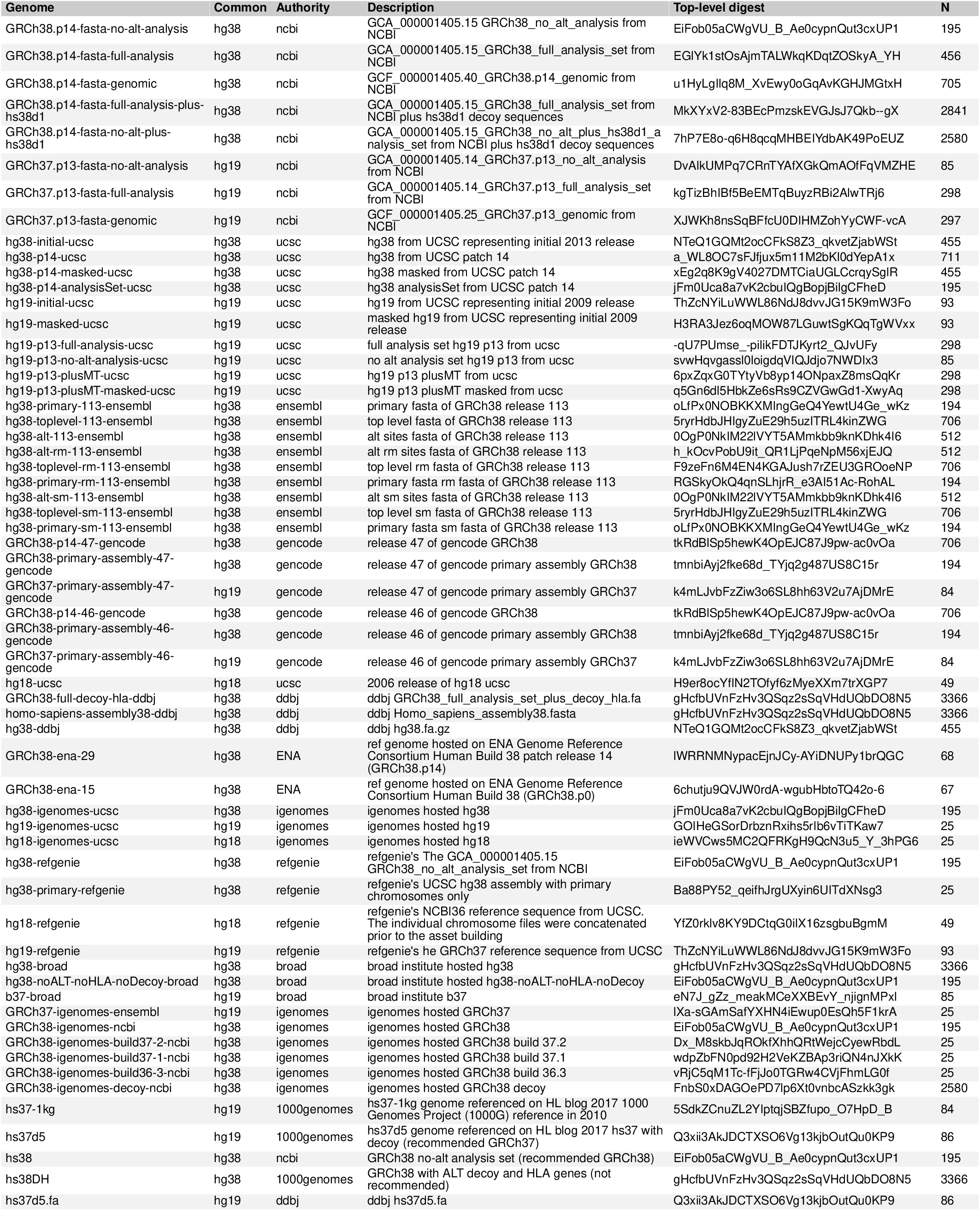
Human reference genomes used for main analysis. N=number of sequences present in the reference genome.

**Supplemental Table S6:**
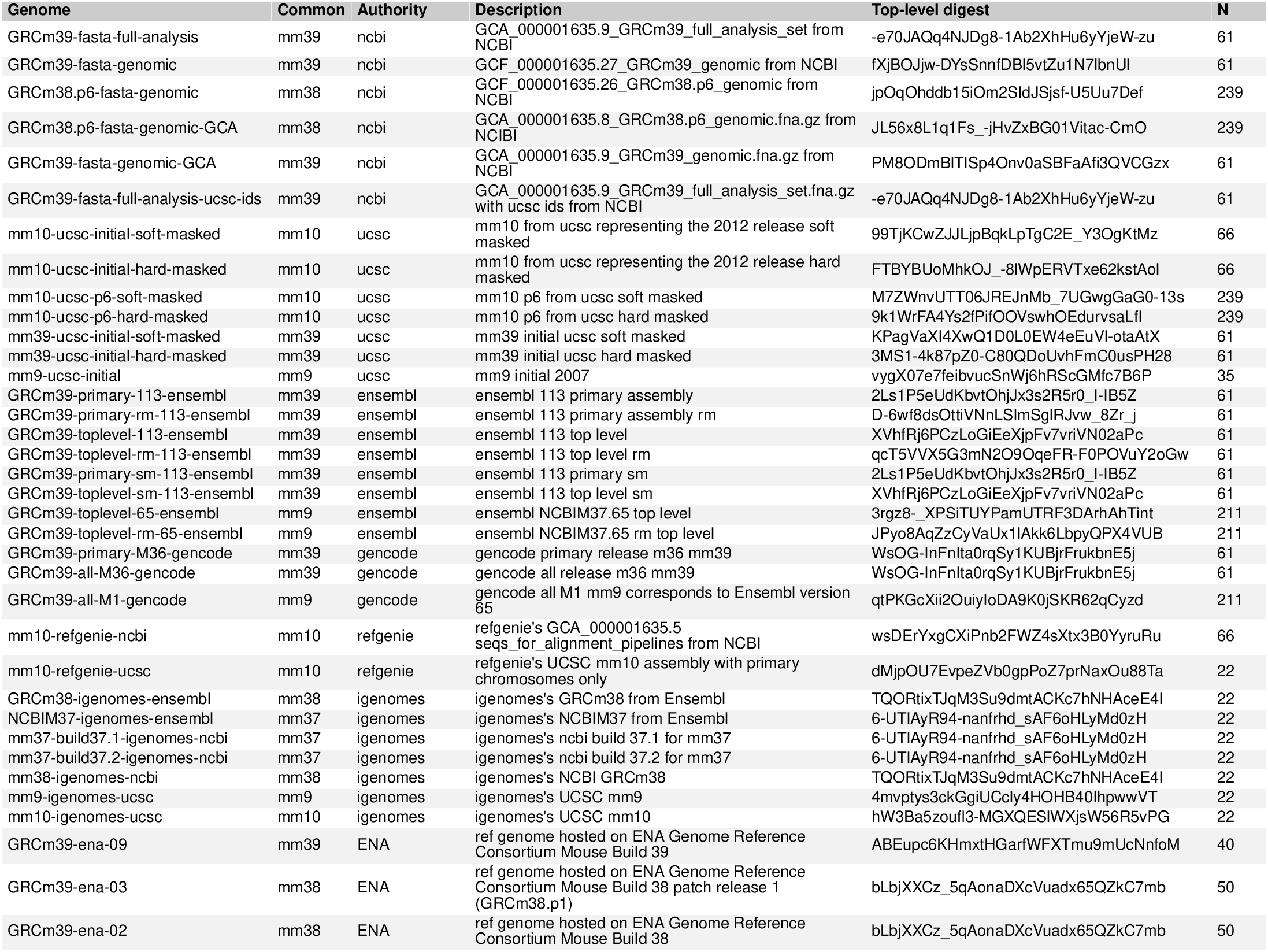
Mouse reference genomes used for main analysis. N=number of sequences present in the reference genome.

